# Genetic inhibition of IL-12β suppresses systolic overload-induced cardiac inflammation and heart failure development

**DOI:** 10.1101/2025.09.05.674485

**Authors:** Umesh Bhattarai, Xiaochen He, Ziru Niu, Lihong Pan, Dongzhi Wang, Hao Wang, Heng Zeng, Jian-Xiong Chen, Joshua S. Speed, John S. Clemmer, John E. Hall, Yingjie Chen

## Abstract

Inflammation promotes heart failure (HF) development, and inhibition of IL-12β simultaneously attenuates interleukin-12 (IL-12) and interleukin-23 (IL-23), two important proinflammatory cytokines. In this study, we used IL-12β knockout (KO) mice to test the hypothesis that genetic inhibition of IL-12β would attenuate transverse aortic constriction (TAC)-induced cardiac inflammation, hypertrophy, and dysfunction, as well as the consequent lung remodeling. IL-12β KO in male and female mice significantly attenuated TAC-induced cardiac dysfunction as evidenced by improved left ventricular (LV) ejection fraction and fractional shortening. IL-12β KO also significantly ameliorated the TAC-induced increase of LV weight, left atrial weight, lung weight, right ventricular (RV) weight, and their ratios to body weight or tibial length in male and female mice. In addition, IL-12β KO significantly attenuated TAC-induced LV leukocyte infiltration, cardiomyocyte hypertrophy, fibrosis, and the consequent lung inflammation and remodeling. Moreover, IL-12β KO reduced TAC-induced alterations of LV gene profile associated with inflammation and fibrosis, as shown by bulk LV RNA sequencing. Furthermore, we found that IL-12β KO significantly attenuated TAC-induced LV accumulation of multiple immune cell subsets, activation of CD4^+^ and CD8^+^ T cells, and the percentage of central memory CD4^+^ and CD8^+^ T cells in the cardiac drainage lymph nodes. Finally, IL-12β KO mice showed significantly reduced IFNγ^+^CD8^+^ and CXCR3^+^CD8^+^ T cells in the drainage lymph nodes as compared with WT after TAC. These findings collectively demonstrate that IL-12β plays a critical role in systolic overload-induced LV inflammation, remodeling, and dysfunction, likely through cardiac immune cell infiltration.

## Introduction

Chronic heart failure (HF), often resulting from left ventricular (LV) failure, is a medical condition in which the LV cannot pump sufficient blood to meet the body’s needs. HF remains a major public health problem and often leads to significant cardiovascular morbidity and mortality (1). HF can result from several different diseases or medical conditions, such as coronary artery disease, systolic hypertension, cardiac valve diseases, diabetes, congenital heart disease, or chronic inflammatory diseases. Even under the current standard clinical care, HF often progresses to WHO class-2 pulmonary hypertension, lung remodeling, and right ventricular (RV) failure (2–6). Patients with HF-induced pulmonary hypertension and RV failure generally have very poor clinical outcomes; thus, identifying novel therapeutic targets for HF treatment is needed.

While the causes or risk factors for HF development are varied, chronic mild or moderate inflammation is often a shared phenomenon among different HF etiologies (7–13). For example, previous studies have demonstrated that pro-inflammatory cytokines such as interleukin-1β (IL-1β) are increased in the heart and circulation of HF patients (7, 8). Also, the Canakinumab Anti-Inflammatory Thrombosis Outcomes Study (CANTOS) trial, which investigated IL-1β blockade with canakinumab (an anti-IL-1β monoclonal antibody) in patients with prior myocardial infarction and evidence of active inflammation, showed reduction in the risk of major cardiovascular events (non-fatal myocardial infarction, non-fatal stroke or death) as compared to standard treatment (9). A subsequent exploratory analysis of the CANTOS data showed that IL-1β inhibition reduced the rate of HF hospitalizations or HF–related death in a dose-dependent manner (10, 11). Moreover, experimental studies from us and others also showed that inhibition of IL-1β signaling significantly attenuated myocardial infarction-induced HF development in rats (12) and transverse aortic constriction (TAC)-induced HF development and/or progression in mice (13).

We and others have also demonstrated that cardiac inflammation and HF development are regulated by immune cell subsets such as CD4^+^ T cells (14), CD8^+^ T cells (15), CD11c^+^ dendritic cells (DCs) (16), NK1.1^+^ cells (17), and macrophages (18) in experimental animals. Macrophages are often the most abundant cardiac immune cells in TAC or myocardial infarction-induced HF animals (18, 19) and a major source of proinflammatory cytokines such as IL-1β, IL-18, IL-12, and IL-23(20). These proinflammatory cytokines could enhance the effective function of other immune cells, such as T cells and NK cells.

IL-12β is an essential subunit for IL-12 and IL-23, two important proinflammatory cytokines mainly produced by activated macrophages and DCs (21–25). Thus, inhibition of IL-12β can simultaneously suppress the proinflammatory effects of IL-12 and IL-23. Since IL-12 promotes IFN-γ production and IL-23 promotes IL-17 production, inhibition of IL-12β simultaneously attenuates the proinflammatory IL-12/IFN-γ and IL-23/IL-17 signaling pathways. Specifically, IL-12 plays an important role in promoting proliferation, activation, and mobilization of CD8^+^ T cells, CD4^+^Th1 cells, and NK cells (21, 26–36), immune cells types that could promote cardiac inflammation and HF secondary to systolic pressure overload produced by TAC, at least in mice (14, 15, 17). Meanwhile, IL-23 plays an important role in stimulating IL-17 production by Th17 cells or γδT17 cells through activating IL-23 receptors expressed on these immune cell subsets through retinoic acid orphan receptor gamma (RORγ)-dependent pathway (23, 24, 37–41). Studies have shown that the IL-23/IL-17 signaling pathway also promotes cardiac inflammation and HF (42). Accordingly, inhibition of IL-12β may be an effective approach to attenuate the IL-12/IFN-γ axis and the IL-23/IL-17 axis, two important pathways that promote HF development.

To test the central hypothesis that IL-12β signaling exerts a critical role in promoting systolic overload-induced cardiac inflammation and HF development, we investigated the role and the underlying mechanisms of IL-12β by using IL-12β knockout (IL-12β KO) mice in TAC-induced LV inflammation and HF.

## Materials and Methods

### Mice and Experimental Protocols

Male and female IL-12β KO (Jackson Lab, Strain #002693) and wild-type (WT) C57BL/6J (Jackson Lab, Strain #000664) mice were used for sham or TAC surgery, a commonly used surgical procedure to mimic clinical conditions such as hypertension or aortic stenosis. Body weight gain/loss was monitored weekly, and LV function was monitored before and every two weeks after TAC. Heart and lung tissue samples were collected at 6 weeks and 4 weeks after TAC for males and females, respectively. Cardiac and pulmonary tissues were harvested and utilized for subsequent histological, immuno-histological, and biochemical analyses. All mice were housed in a temperature-controlled environment with 12-hour light/dark cycles. This study was approved by the Animal Care and Use Committee at the University of Mississippi Medical Center.

### TAC Procedure

TAC surgery was performed using a 26-gauge and 27-gauge blunt needle for males and females, respectively, after anesthetizing the mice with an intraperitoneal injection of ketamine (100 mg/Kg) and xylazine as previously described (10 mg/Kg) (43, 44).

### Echocardiography

Echocardiography was performed using a VisualSonics Vevo 3100 imaging system (FUJIFILM VisualSonics Inc., Canada) as previously described (44). Briefly, the mice were anesthetized by inhalation of 1-2% isoflurane mixed with 100% oxygen. M-mode echocardiographic images were taken and analyzed using Vevo LAB software (FUJIFILM VisualSonics Inc., Canada) to measure heart rate, LV ejection fraction, LV fractional shortening, LV end-systolic diameter, LV end-diastolic diameter, LV end-systolic volume, LV end-diastolic volume, LV anterior and posterior wall thickness at end-systole or end-diastole, stroke volume, and cardiac output. The pressure gradient was assessed using the blood flow velocity difference across the TAC site by the pulsed-wave Doppler method.

### Histological Staining

The histological and immunological staining were performed according to previous studies (45). The detailed method for histological and immune staining is also provided in the supplemental data.

### Flow Cytometry Analyses

LV tissues were harvested, minced into small pieces, and digested in HBSS supplied with Deoxyribonuclease I (66.7 μg/mL, Sigma-Aldrich) and LiberaseTM (125 μg/mL, Roche Diagnostics, Germany) at 37°C for 30 minutes using a tissue dissociator (Miltenyi Biotec), followed with cell purification and immune staining as we previously described(17, 46). Data were acquired on a BD FACSymphony^TM^ A3 Cell Analyzer (BD Biosciences) and analyzed by using FlowJo-v10 (FlowJo, OR) software. The gating strategies used for LV flow cytometry analysis are presented in Supplementary Figure 1.

For the flow cytometry from lymph nodes, the lymph nodes were mechanically dissociated by gently pressing them with the plunger of a syringe and filtered through a 100 µm cell strainer. The staining procedure is the same as described above. The gating strategies used for lymph node flow cytometry analysis are presented in Supplementary Figure 2. For cytokine production assay, the isolated immune cells were stimulated with 1X Cell Stimulation Cocktail (Invitrogen, 00-4970-93) and 1X Protein Transport Inhibitor Cocktail (00-4980-93) in RPMI 1640 with 10% FBS at 37°C for 2 hours as we previously described(17, 46). The cells were then stained with antibodies for cell surface markers, permeabilized, and stained with antibodies against different intracellular cytokines (Supplementary Table 1).

### Western Blot Analyses

Western blot analysis was performed as previously described (45), and the detailed method is provided in the supplemental data.

### Statistical Analysis

Data are presented as Mean±SEM. A two-way ANOVA followed by a Bonferroni post-hoc test was used to determine statistical differences between WT and IL-12β KO mice under sham or TAC conditions. An unpaired t-test was used to test statistical differences between the 2 groups. The log-rank test was used to analyze survival curves. All the statistical tests were performed using GraphPad Prism 10 software. p<0.05 was considered statistically significant.

## Results

### IL-12β KO significantly attenuated TAC-induced LV dysfunction in male and female mice

To determine whether cardiac IL-12β expression is changed during TAC-induced HF, a Western blot was performed in LV tissues from mice after sham and TAC surgery. Our results show that LV IL-12β protein expression was significantly increased in WT mice after TAC (Fig. 1A). To determine the role of IL-12β in TAC-induced cardiac inflammation and HF, IL-12β KO mice and WT mice of both sexes were used for sham or TAC surgery. IL-12β KO significantly attenuated TAC-induced mortality rate in female mice (1 out of 23) as compared with female WT mice (15 out of 31). TAC-induced mortality was not statistically different between male IL-12β KO (5 out of 19) mice and male WT mice (7 out of 17) (Fig. 1B). The blood flow velocity difference across the TAC site was similar between WT TAC mice and IL-12β KO TAC mice (Supplementary Figure 3), indicating comparable TAC between IL-12β KO and WT mice. Echocardiography further showed that LV function and LV dimensions were comparable between WT and IL-12β KO mice under sham conditions. However, as compared with corresponding WT mice, IL-12β KO significantly attenuated TAC-induced LV dysfunction and LV dilatation in male and female mice (Fig. 1C and Supplementary Figure 4).

**Figure 1:**
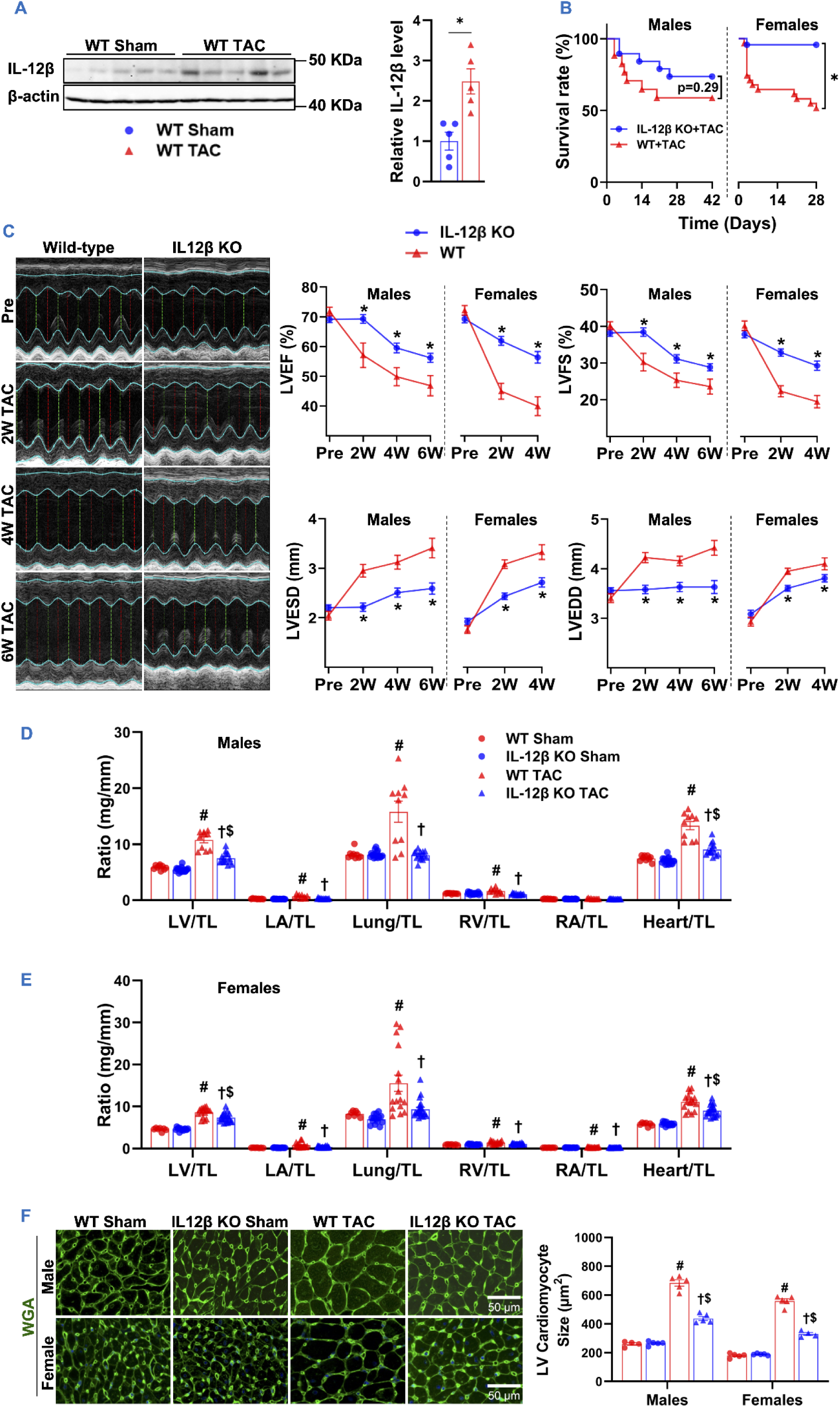
IL-12β KO significantly attenuated TAC-induced cardiac dysfunction, an increase in LV weight, LA weight, lung weight, and RV weight in male and female mice. (A) Western blots and quantification of LV IL-12β expression in WT sham and WT TAC mice. (B) Survival curves of WT TAC and IL-12β KO TAC mice of both sexes (log-rank test). (C) Representative M-mode echocardiographic images of WT and IL-12β KO male mice: pre-TAC, 2 weeks, 4 weeks, and 6 weeks after TAC, and Quantified data of echocardiographic measurements of LVEF, LVFS, LVESD, and LVEDD of both sexes. (D, E) The ratio of LV weight, left atrial (LA) weight, lung weight, RV weight, right atrial (RA) weight, and total heart weight to tibial length (TL) of the indicated groups. (F) Representative LV WGA staining images and quantified data of cardiomyocyte cross-sectional areas. *p<0.05; ^#^p<0.05 compared to WT sham; ^†^p<0.05 compared to WT TAC; ^$^p<0.05 compared to IL-12β KO sham; n = 7 to 22 per group; LVEF, LV ejection fraction; LVFS, LV fractional shortening; LVESD, LV end-systolic diameter; LVEDD, LV end-diastolic diameter.

### IL-12β KO significantly attenuated TAC-induced LV hypertrophy, increases in lung weight, and LV myocyte hypertrophy in male and female mice

LV weight, left atrial (LA) weight, lung weight, RV weight, and their ratios to tibial length or body weight were comparable between WT and IL-12β KO mice under sham conditions (Fig. 1D-E, Supplementary Tables 2 and 3, and Supplementary Figure 4). TAC significantly increased LV weight, LA weight, lung weight, RV weight, and their ratios to tibial length or body weight in male and female WT mice, but these changes were significantly attenuated in IL-12β KO mice as compared with corresponding WT mice (Fig. 1D-E, Supplementary Tables 2 and 3, and Supplementary Figure 4). Moreover, wheat germ agglutinin (WGA) staining was performed to measure LV cardiomyocyte cross-sectional area. The data showed that the TAC-induced increase in LV cardiomyocyte cross-sectional area was significantly reduced in IL-12β KO mice (Fig. 1F).

### IL-12β KO protected hearts against TAC-induced changes in the LV gene profile

To determine the underlying mechanisms of IL-12β KO in protecting mice from TAC-induced LV hypertrophy and dysfunction, bulk RNA sequencing was performed in LV tissues from female WT and IL-12β KO mice (Fig. 2). A principal component analysis (PCA) of LV RNA-seq data revealed distinct clusters of LV gene profiles among the WT TAC group, IL-12β KO TAC group, and the sham groups (both WT and IL-12β KO mice). PCA also showed that sham WT and IL-12β KO mice are clustered together, while IL-12β KO TAC mice were clustered near the sham mice (Fig. 2A).

**Figure 2:**
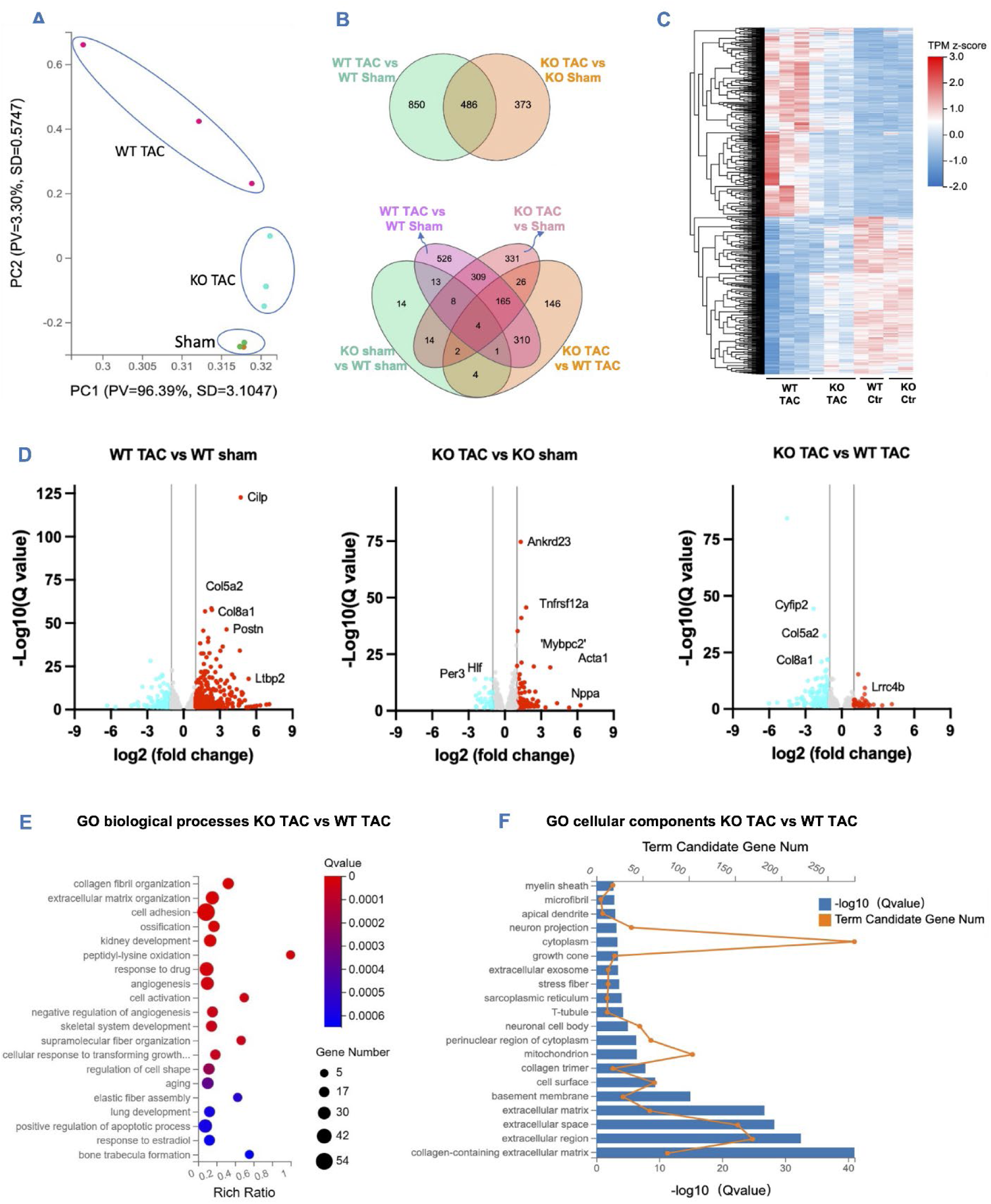
IL-12β KO attenuated TAC-induced alterations in the gene expression related to fibrosis and inflammation. (A) Principal component analysis of WT and IL-12β KO mice, under sham or TAC conditions. (B) Venn diagram showing differentially expressed genes (DEGs) of WT and IL-12β KO mice under sham or TAC conditions, as well as the shared and uniquely changed genes. (C) DEGs cluster heatmap in WT mice compared to IL-12β KO mice, under sham or TAC conditions. (D) Volcano plots showing upregulated and downregulated genes among the groups. (E) Gene ontology (GO) biological process enrichment bubble chart in WT TAC mice compared to IL-12β KO TAC. (F) GO cellular component enrichment histogram shows the enriched cellular components between KO and WT mice after TAC. n = 2/group for sham and n=3/group for TAC.

We further determined the differentially expressed genes (DEGs) among the experimental groups. Under basal/sham conditions, there were only 60 DEGs (39 upregulated and 21 downregulated) in LV tissues between IL-12β KO mice and WT mice (Supplementary Figure 5). As compared with corresponding control mice, TAC caused 1336 DEGs in WT mice (728 upregulated vs 608 downregulated), and 859 DEGs in IL-12β KO mice (395 upregulated vs 464 downregulated) (Supplementary Figure 5). Comparison of LV gene profiles between IL-12β KO TAC and WT TAC mice revealed 658 DEGs (232 upregulated vs 426 downregulated) (Supplementary Figure 5). Venn diagrams were generated showing the DEGs of WT and IL-12β KO mice after TAC as compared with their corresponding sham groups, as well as the uniquely expressed and shared DEGs among these four experimental groups (Fig. 2B).

A heatmap was also generated for the four experimental groups using the DEGs between WT and IL-12β KO mice after TAC (Fig. 2C). The results clearly showed that IL-12β KO drastically attenuated TAC-induced alterations of LV gene profiles as LV gene profiles of IL-12β KO mice are like the sham mice (Fig. 2C). Moreover, the DEGs that changed over 2-fold among these groups are also presented with volcano plots (Fig. 2D). Together, these findings clearly demonstrate that IL-12β KO mice are protected from TAC-induced LV remodeling.

### IL-12β KO suppressed LV genetic pathways associated with extracellular matrix remodeling, inflammation, and LV fibrosis in mice after TAC

To reveal the major molecular signaling pathways changed between WT and KO mice after TAC, pathway analyses were further performed for LV DEGs of WT and IL-12β KO mice after TAC. The gene ontology (GO) biological process enrichment analysis showed that the top enriched pathways were collagen fibril organization, extracellular matrix organization, cell adhesion, ossification, angiogenesis, and TGFβ signaling for DEGs between WT and KO mice after TAC (Fig. 2E). The GO cellular component enrichment analysis showed that extensively enriched cellular components were collagen-containing extracellular matrix, extracellular region, extracellular matrix, and other extracellular matrix-related cellular components (Fig. 2F). The top enriched GO molecular pathways were extracellular matrix structural constituent, heparin-binding, integrin binding, and protein binding, suggesting that IL-12β deficiency is associated with major impact on extracellular matrix organization and fibrosis, as well as inflammation under pressure overload conditions (Supplementary Figure 5). Kyoto Encyclopedia of Genes and Genomes (KEGG) enrichment analysis of LV DEGs shows the top enriched pathways are protein digestion and absorption, diabetic cardiomyopathy, AGE-RAGE signaling in diabetic complications, and ECM-receptor interaction pathways (Supplementary Figure 5).

To further determine statistically upregulated and downregulated functions/pathways for DEGs between the WT TAC group and the IL-12β KO TAC group, Gene Set Enrichment Analysis (GSEA) with ReactomeGSA was further performed for DEGs of the WT TAC group and the IL-12β KO TAC group. The findings showed that the top downregulated functions/pathways of IL-12β KO and WT mice after TAC are collagen formation, extracellular matrix organization, collagen biosynthesis and modifying enzymes, assembly of collagen fibrins and other multimeric structure, collagen chain trimerization etc., as well as pathways related to immune response such as integrin cell surface integrations, *immunomodulatory interactions between lymphatic and non-lymphatic cells*, and DAP12 interactions [DAP12 is an immunoreceptor tyrosine-based activation motif (ITAM)-bearing adapter molecule that transduces activating signals in NK and myeloid cells]. (Fig. 3A, B).

**Figure 3.**
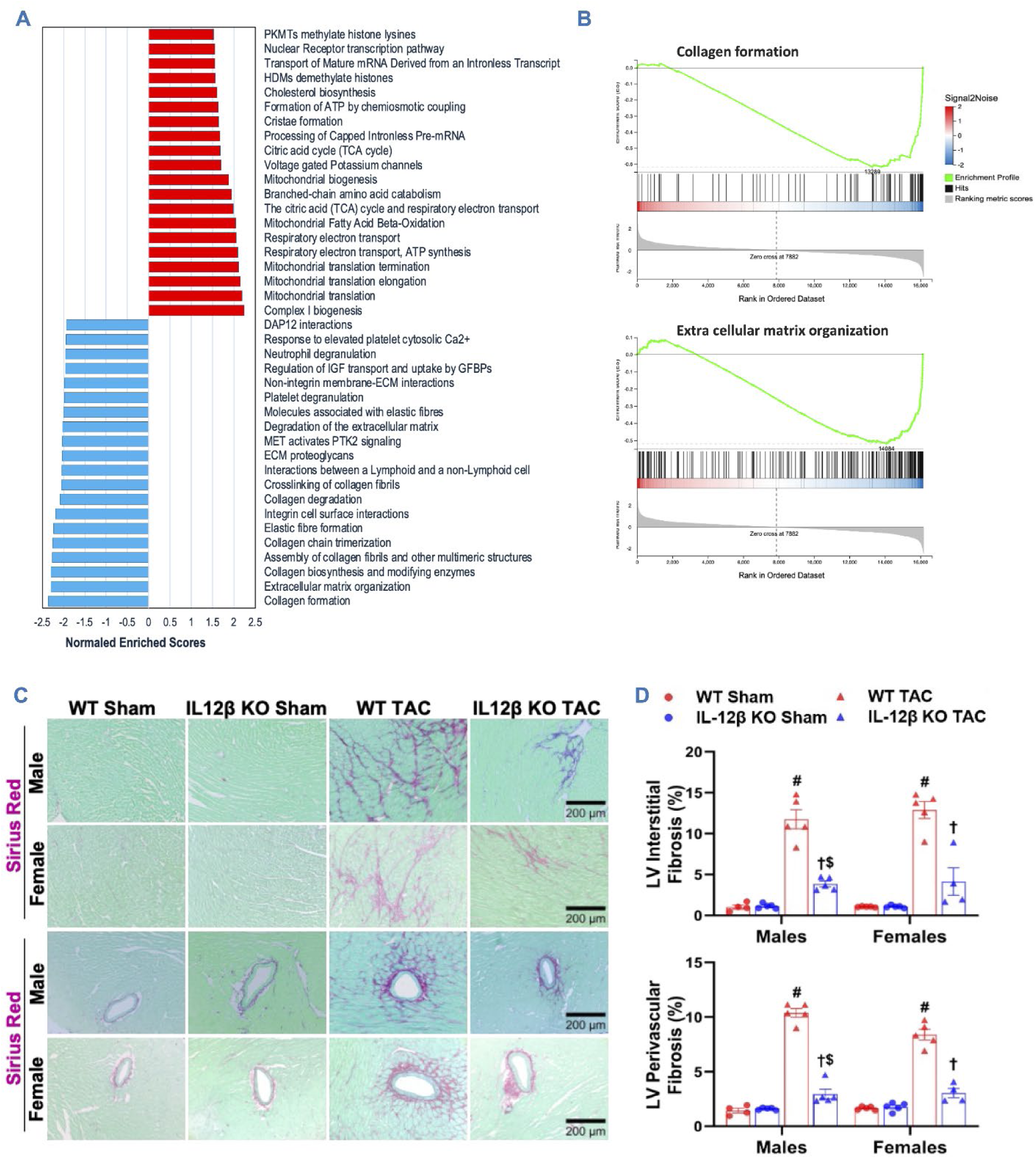
IL-12β KO suppressed LV genetic pathways associated with extracellular matrix remodeling, inflammation, and LV fibrosis in mice after TAC. (A) The top 20 upregulated and downregulated pathways between KO and WT mice after TAC, identified by GSEA with ReactomeGSA. (B) Representative GESA plots of collagen formation and extracellular matrix organization. (C, D) Representative images and quantified data of LV interstitial and perivascular fibrosis performed by Sirius Red/Fast Green staining in male and female mice. ^#^p<0.05 compared to WT sham; ^†^p<0.05 compared to WT TAC; ^$^p<0.05 compared to IL-12β KO sham; n = 4-5 per group.

ReactomeGSA also showed that the top upregulated functions/pathways include mitochondrial complex I biogenesis, mitochondrial translation, mitochondrial translation elongation, mitochondrial fatty acid beta oxidation, and branched-chain amino acid catabolism (Fig. 3A), indicating that IL-12β KO mice were protected from TAC-induced LV mitochondrial and metabolic dysfunction in mice. Consistent with the findings that IL-12β KO suppressed LV genetic pathways associated with extracellular matrix remodeling, histological staining showed that IL-12β KO significantly attenuated TAC-induced increase in LV interstitial and perivascular fibrosis (Fig. 3C, D). Overall, GSEA by ReactomeGSA and KEGG pathway showed that IL-12β KO significantly downregulated LV pathways associated with extracellular matrix processing, immune and inflammatory responses, and the interactions between immune cells with nonimmune cells to reduce inflammation. IL-12β KO also upregulated LV pathways, enhancing mitochondrial biogenesis and metabolism in mice after TAC.

### IL-12β KO suppressed TAC-induced LV genetic pathways associated with inflammation and LV leukocyte infiltration in mice

Since GSEA shows that IL-12β KO significantly suppressed TAC-induced LV genetic pathways associated with inflammatory responses and the interactions between immune cells and nonimmune cells (Fig. 3A), GSEA by KEGG pathway was also performed. The findings showed that downregulated pathways are associated with various pathogen infection (such as pathways Staphylococcus aureus infection, Malaria infection, Human papillomavirus infection, EB virus infection etc.), immune or inflammatory responses (such as ECM-receptor interaction, Phagosome, Complement and coagulation cascades, Lysosome, Focal adhesion, Chronic myeloid leukemia, Rheumatoid arthritis, and Antigen processing and presentation etc.), and metabolic pathways associated cell recruiting and injury repair (such as Protein digestion and absorption, Focal adhesion, Glycosaminoglycan biosynthesis, TGF-beta signaling pathway etc.) (Fig. 4A).

**Figure 4:**
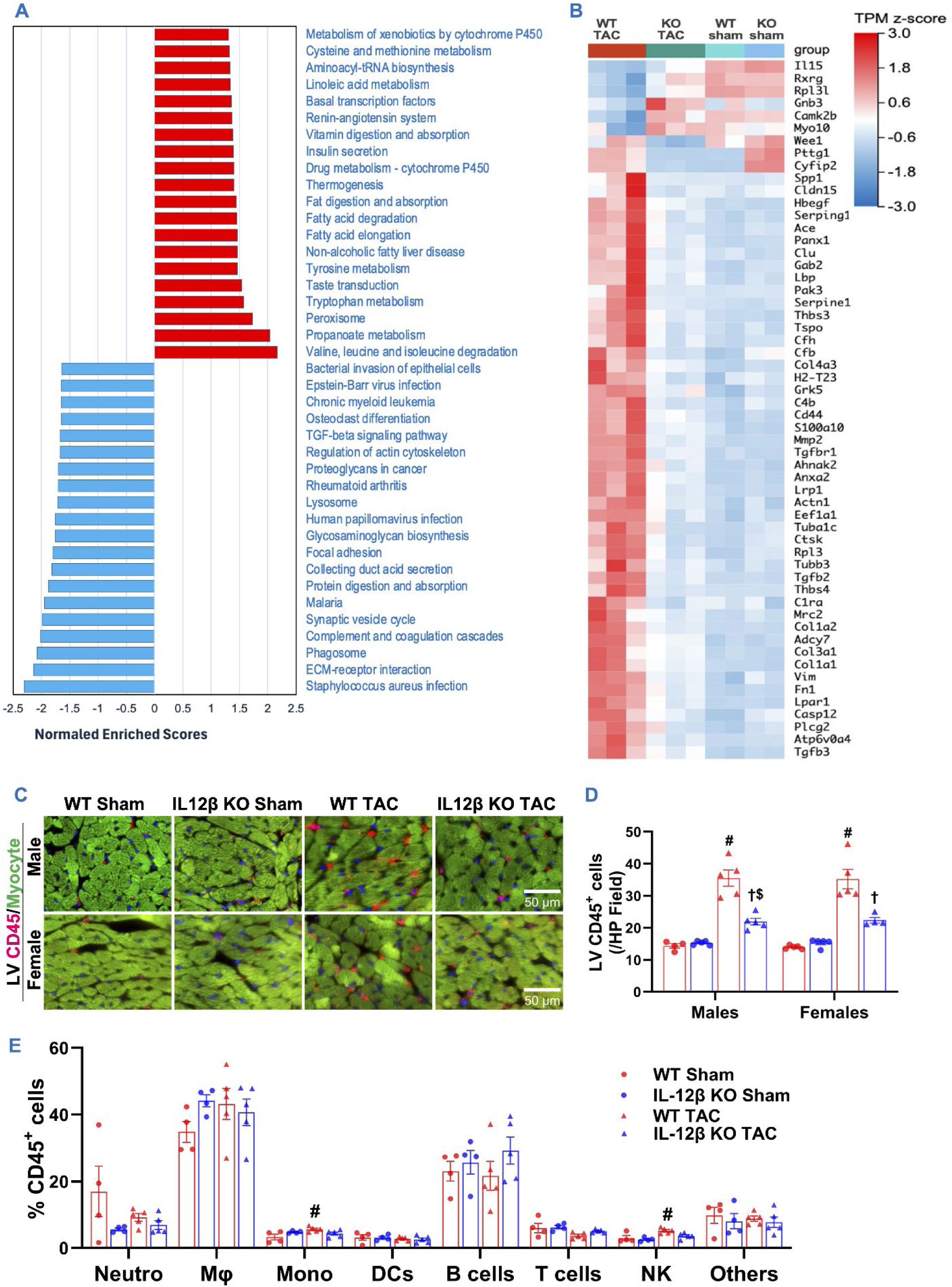
IL-12β KO significantly attenuated TAC-induced LV inflammation in mice. (A) The top 20 upregulated and downregulated pathways between KO and WT mice after TAC identified by GSEA with KEGG pathway analysis. (B) Heatmap shows the top LV immune and/or inflammation-related genes in KO and WT mice after TAC. (C, D) Representative LV CD45^+^ immuno-staining images and quantified data of LV CD45^+^ of the indicated groups. (E) Percentage distribution of major immune cell subsets within LV CD45^+^ cells of the indicated groups determined by flow cytometry. ^#^p<0.05 compared to WT sham; ^†^p<0.05 compared to WT TAC; ^$^p<0.05 compared to IL-12β KO sham; n = 4-5 per group.

We further analyzed DEGs associated with infection, immune diseases, and the immune system of WT and IL-12β KO mice after TAC based on the KEGG Pathway Classification analysis (Supplementary Figure 6). We found a total of 105 DEGs, and the top upregulated genes include Gnb3, Camk2b, Gadd45a, IL15, Rpl3l, Camk2a, Bcl2L11 etc., and the top downregulated DEGs include Cyfip2, Col1a2, Vim, Mmp2, Tgfb2, Fn1, C4b, Ace, Thbs4, and Spp1. To reveal the relative expressions of these immune-related DEGs among the four experimental groups, a heatmap was generated for the top 56 DEGs (p<0.01, and net Log2 fold change is greater than 0.5) (Figure 4B). The top downregulated immune related genes DEGs include adhesion molecules recruiting immune cells (such as Cxadr, CD44, Icam1, Itgb1 or CD29), complements, a group of genes modulating the interactions between CD8 and other cells (B2m, and MHCI molecules such as H2-T23, H2-K1, H2-Q7), and a group of collagens (such as Col1a2, Col1a1, and Col3a1) that are implicated in the development of inflammation (Figure 4B). These findings suggest that cardiac immune signaling pathways related to cardiac immune cell infiltration, particularly the signaling related to CD8 T cell activity, are reduced in IL-12β KO mice after TAC.

We further determined the effect of IL-12β KO on TAC-induced LV immune cell infiltration by using immuno-histology and flow cytometry. IL-12β KO had no detectable effects on LV CD45^+^ leukocyte infiltration under sham conditions, but IL-12β KO significantly attenuated TAC-induced LV leukocyte infiltration in male and female mice (Fig. 4C, D). However, to our surprise, the percentages of the major immune cell subsets within LV CD45^+^ cells (such as macrophages, B cells, T cells, and neutrophils) were largely unaffected in IL-12β KO mice after sham or TAC (Fig. 4E).

### IL-12β KO significantly attenuated TAC-induced pulmonary immune cell infiltration, fibrosis, and vessel remodeling in male and female mice

Our previous studies demonstrated that LV failure causes pulmonary remodeling and the consequent RV hypertrophy (2, 13, 47). Thus, we further determined the effect of IL-12β KO on TAC-induced pulmonary remodeling. Although IL-12β KO had no detectable effects on pulmonary CD45^+^ leukocyte infiltration, fibrosis, and arteriole muscularization as compared to WT mice under sham conditions (Supplementary Figure 7), IL-12β KO significantly attenuated TAC-induced pulmonary CD45^+^ leukocyte infiltration, fibrosis, and vessel muscularization in both male and female mice (Supplementary Figure 7). These findings indicate that IL-12β KO also significantly attenuated TAC-induced HF progression in mice of both sexes.

### IL-12β KO significantly attenuated TAC-induced early-phase LV dysfunction, cardiomyocyte hypertrophy, inflammation, and fibrosis

TAC-induced LV immune cell infiltration often peaks ∼7 days after TAC, a time point at which pulmonary remodeling is not observed. To understand the impact of IL-12β on early-stage LV inflammation and remodeling, we further determined the cardiac immune cell infiltration and the immune cell subsets in WT and IL-12β KO mice 7 days after TAC. Interestingly, IL-12β KO already significantly attenuated TAC-induced LV dysfunction 7 days after TAC (Fig. 5A). The TAC-induced increases in LV end-systolic diameter (LVESD) and end-systolic volume (LVESV) were significantly reduced in IL-12β KO mice as compared to WT mice, while LV end-diastolic diameter (LVEDD) and LV end-diastolic volume (LVEDV) were similar between WT and IL-12β KO mice (Fig. 5A and Supplementary Figure 8). In addition, TAC-induced LV hypertrophy, LV cardiomyocyte hypertrophy, LV CD45^+^ infiltration, and LV fibrosis were significantly decreased in IL-12β KO mice as compared to WT mice (Fig. 5B-D and Supplementary Figure 8).

**Figure 5:**
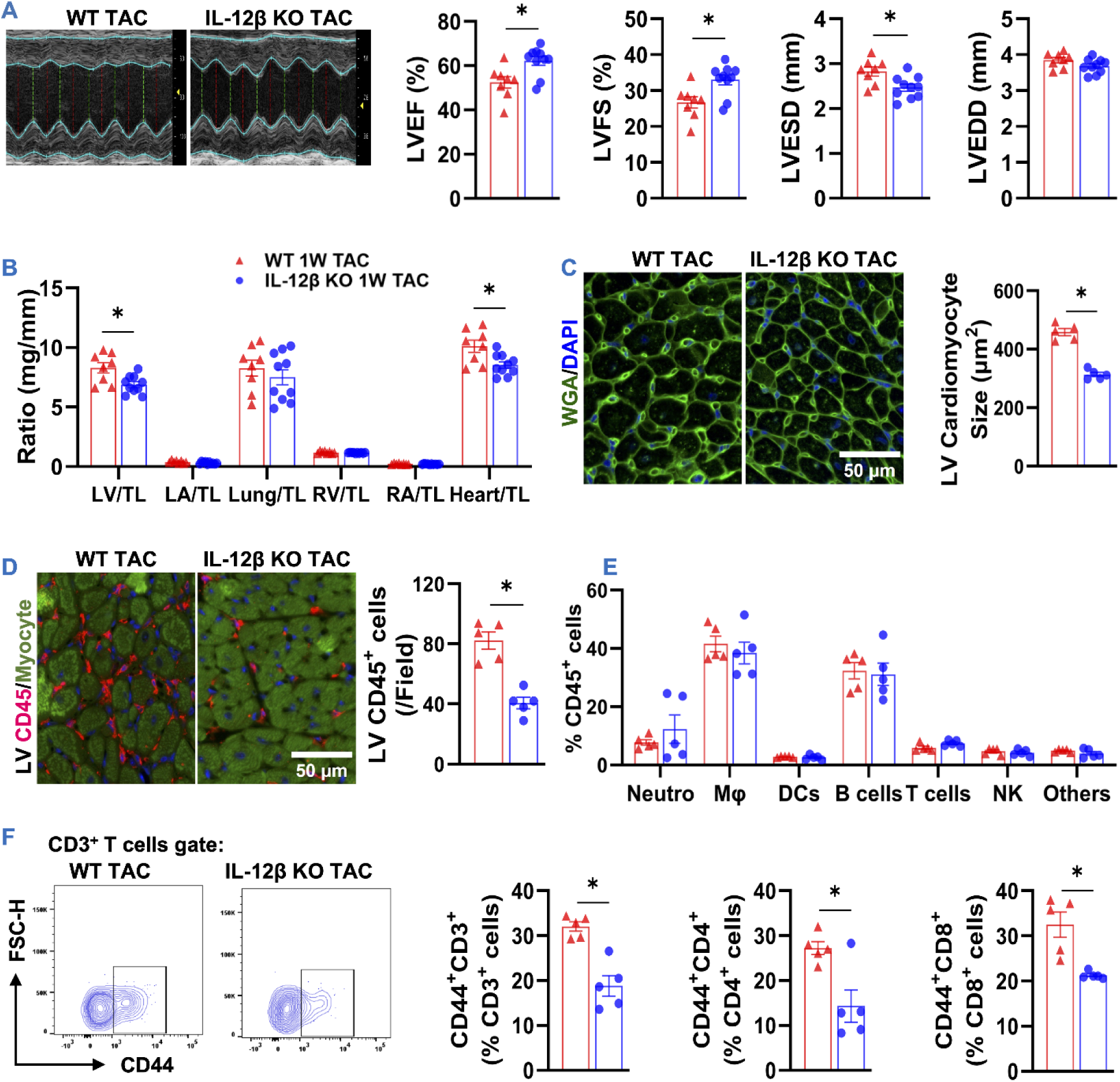
IL-12β KO significantly attenuated TAC-induced early-phase LV dysfunction, cardiomyocyte hypertrophy, and inflammation. (A) Representative M-mode echocardiographic images and quantified data of echocardiographic measurements of LVEF, LVFS, LVESD, and LVEDD of the indicated groups. (B) The ratios of cardiac and pulmonary tissues to tibial length (TL) of the indicated groups. (C, D) Representative images and quantified data of LV cardiomyocyte size and LV CD45^+^ cells of the indicated groups. (E) Percentage of major immune cell subsets within LV CD45^+^ cells determined by flow cytometry. (F) Representative flow cytometry plots and quantified data of CD44^+^CD3^+^, CD44^+^CD4^+^, and CD44^+^CD8^+^ T cells within the corresponding T cells. *p<0.05; n = 5-10 per group.

We further performed flow cytometry to detect the major LV immune cell subsets. Again, the percentages of LV macrophages, B cells, T cells, and neutrophils were similar between WT and IL-12β KO mice 7 days after TAC (Fig. 5E). Given the marked increase in LV CD45^+^ cell infiltration in WT mice compared to IL-12β KO mice (Fig. 5C), it can be inferred that the major immune cell subsets increased proportionally following TAC, and this response was attenuated in IL-12β KO mice. CD44 plays an important role in tissue recruiting immune cells and stem cells, and CD44 expression is often used as a marker of T cell activation. Interestingly, the percentages of LV CD44^+^CD3^+^ T cells, CD44^+^CD4^+^ T cells, and CD44^+^CD8^+^ T cells were significantly attenuated in IL-12β KO mice as compared to WT mice after TAC (Fig. 5F). The average CD44 expression in CD4^+^ T cells and CD8^+^ T cells were also significantly reduced in IL-12β KO mice after TAC. These findings indicate that IL-12β KO significantly attenuated cardiac dysfunction and inflammation as early as 7 days after TAC.

### IL-12β attenuated “stem cell-like” central memory T cell activation and IFNγ^+^CD8^+^ T cells in cardiac drainage lymph nodes

Up to now, we found that IL-12β KO significantly suppressed TAC-induced cardiac immune cell infiltration and T cell activation, while the LV gene profile also showed that IL-12β KO significantly suppressed cardiac immunomodulatory interactions between lymphatic and non-lymphatic cells. Since the drainage lymph nodes play a crucial role in immune surveillance by promoting T cell proliferation, activation, and enhancing effective function by increasing their tissue homing capacity and production of cytotoxic molecules such as IFNγ. Since TAC-induced cardiac inflammation generally peaks ∼7 days after TAC, a time point at which there is no detectable effect on pulmonary immune cell infiltration and dysfunction, we investigated the immune cell composition and their activation in cardiac drainage lymph nodes. Surprisingly, we found that the percentage of major immune cell subsets within CD45^+^ cells and the percentages of major T cell subsets within CD3^+^ T cells were similar between WT and IL-12β KO mice under control conditions and after TAC (Fig 6A, B). IL-12β KO also had no detectable effect on the percentages of Tregs and IL10^+^Tregs (data not shown) in the drainage lymph nodes in mice under control conditions and after TAC. However, we found that IL-12β KO abolished TAC-induced increase of IFNγ^+^CD8^+^ cells without affecting the percentage of TNFα^+^CD8^+^ T cells (Fig. 6C), while the percentages of IFNγ^+^CD4^+^ T cells and TNFα^+^CD4^+^ T cells were similar between KO and WT mice under control conditions or after TAC (Supplementary Figure 9). Since CD8^+^ T cells generally have higher expression of chemokine receptor CXCR3, CXCR3 plays an important role in promoting the CD8^+^ T cells’ infiltration into injured tissues. Interestingly, the percentages of CXCR3^+^CD8^+^ cells and CXCR3^+^CD4^+^ T cells in the drainage lymph nodes were significantly reduced in IL-12β KO after TAC as compared with corresponding WT mice (Fig. 6D, E).

**Figure 6:**
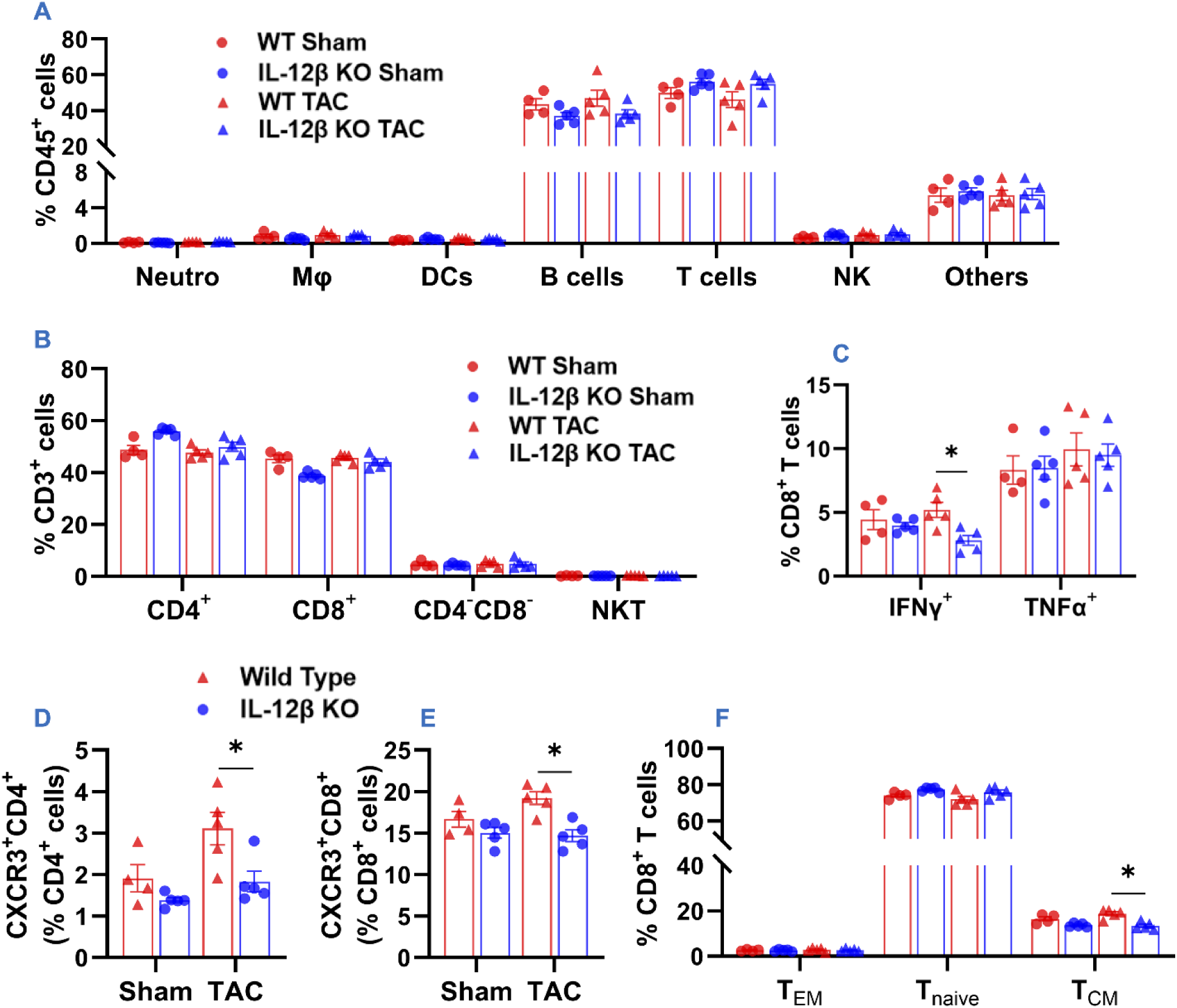
IL-12β KO significantly attenuated “stem cell-like” central memory T cell activation and their IFNγ production in the drainage lymph nodes. (A, B) Percentages of major immune cell subsets within CD45^+^ cells and the T cell subsets within CD3^+^ cells in the drainage lymph nodes determined by flow cytometry. (C) Percentages of IFNγ^+^CD8^+^ and TNFα^+^CD8^+^ within CD8^+^ T cells. (D, E) Percentages of CXCR3^+^CD4^+^ and CXCR3^+^CD8^+^ T cells within CD4^+^ and CD8^+^ T cells, respectively. (F) Percentage of effective memory (CD44^+^CD62L^-^), naïve (CD44^-^CD62L^+^), and central memory (CD44^+^CD62L^+^) T cells of CD8^+^ T cells. *p<0.05; T_EM_, Effective Memory T Cells; T_naïve_, Naïve T Cells; T_CM_, Central Memory T Cells; n = 5 per group.

“Stem cell-like” central Memory T cells (CMTs, generally characterized as CD44^+^CD62^+^ T cells) primarily reside in lymphatic tissues such as drainage lymph nodes. CMTs are characterized by their ability to rapidly proliferate and differentiate into effector T cells upon re-exposure to APC-presented antigens. Interestingly, TAC caused significant increases in the percentages of CD4^+^CD44^+^CD62^+^ and CD8^+^CD44^+^CD62^+^ central memory T cells within their corresponding T cell subsets in WT mice, while IL-12β KO abolished TAC-induced increases of these central memory T cells (Fig. 6F and Supplementary Figure 9). Moreover, CD44 expression in both CD4^+^ and CD8^+^ T cells was significantly reduced in KO mice as compared with WT mice after TAC (data not shown).

## Discussion

We tested the hypothesis that genetic inhibition of IL-12β would attenuate pressure overload-induced HF development and progression by using WT and IL-12β-deficient mice. There are several major new findings. First, the expression of IL-12β in LV tissue was significantly increased in WT TAC mice as compared to WT sham mice. Second, IL-12β KO significantly attenuated TAC-induced mortality in male mice as well as LV hypertrophy and dysfunction in male and female mice. Third, bulk RNA sequencing of LV tissues showed that IL-12β KO protected hearts against TAC-induced gene profile alterations and pathways related to extracellular collagen formation, organization, and remodeling. Fourth, bulk RNA sequencing of LV tissues also showed that IL-12β KO significantly attenuated TAC-induced LV upregulations of genes and signaling pathways promoting inflammatory responses, immune cell recruitments, and the interactions between lymphatic cells and non-lymphatic cells. Fifth, IL-12β KO significantly attenuated TAC-induced LV inflammation and fibrosis, consequent lung inflammation and remodeling, and RV hypertrophy. Finally, we found that IL-12β KO significantly attenuated TAC-induced cardiac dysfunction and inflammation as early as 7 days after TAC. This was associated with significantly reduced LV T cell activation, central memory T cell activation, and IFNγ^+^CD8^+^ cells in cardiac drainage lymph nodes.

One of the major findings of this study is that IL-12β KO significantly attenuated TAC-induced cardiac dysfunction in male and female mice, as evidenced by the following changes. First, IL-12β KO significantly attenuated the TAC-induced decrease of LV ejection fraction, LV fractional shortening, and increase of LV end-systolic and end-diastolic diameter and volume. In addition, IL-12β KO significantly ameliorated TAC-induced LV inflammation, hypertrophy, and fibrosis. Moreover, the LV gene profile also demonstrated that IL-12β KO effectively protected the heart against TAC-induced LV remodeling.

The findings of significant cardiac dysfunction, LV inflammation, hypertrophy, and fibrosis in WT mice after TAC are consistent with the notion that inflammation plays an important role in pressure overload-induced HF development. Previous studies from us and others have demonstrated the important role of immune cells such as CD4^+^ T cells(14), CD8^+^ T cells(15), CD11c^+^ dendritic cells(16), and NK1.1^+^ cells(17) in pressure overload-induced cardiac inflammation and dysfunction. The finding that IL-12β KO significantly attenuated TAC-induced HF not only reinforces the critical role of inflammation in HF development(2, 48) but also highlights the specific contribution of IL-12β to cardiac inflammation and HF development. However, all the above parameters were comparable between WT and IL-12β KO mice under sham conditions, suggesting that IL-12β deficiency does not significantly impact cardiac structure and function under unstressed conditions.

In addition, IL-12β KO significantly attenuated the TAC-induced increase in lung weight, RV weight, and their ratio to tibial length or body weight, indicating that IL-12β not only plays a critical role in pressure overload-induced HF development, but also in HF progression. This is further supported by findings showing that IL-12β KO significantly reduced pulmonary inflammation, fibrosis, and arteriole muscularization. Since LV failure could contribute to pulmonary inflammation, structural remodeling, and RV hypertrophy, the reduced lung inflammation, remodeling, and RV hypertrophy in IL-12β KO mice after TAC could be partially due to improved cardiac function. However, given that lung inflammation can drive pulmonary remodeling and RV hypertrophy independently of LV function in mice with pre-existing HF (47), IL-12β deficiency could directly attenuate lung inflammation and remodeling in HF mice. Inflammation not only contributes to the onset of HF but also plays a crucial role in its progression (2, 47, 48). We have already demonstrated that HF is associated with a substantial accumulation and activation of macrophages, CD11c^+^ dendritic cells, and T cell activation in the lungs, while modulating inflammatory response effectively attenuated chronic HF-induced pulmonary inflammation, remodeling, and RV hypertrophy in mice with pre-existing HF(13, 47, 49). For example, aggravating lung inflammation by exposing mice to PM2.5 worsened HF-induced lung inflammation and remodeling in mice with pre-existing HF(46), whereas promoting endogenous induction of regulatory T cells (Tregs)(48) or inhibiting IL-1β(13) attenuated HF progression in mice with pre-existing HF. These findings support the concept that inflammation not only plays a critical role in pressure overload-induced HF development but also in HF progression(16, 48, 49).

Another important finding of our study is that IL-12β KO suppressed TAC-induced LV gene expression pathways associated with inflammation and LV leukocyte infiltration in mice. HF-induced cardiac dysfunction in WT mice is also corroborated by bulk RNA sequencing data from LV tissues showing significant upregulation of genes and enrichment of many pathways related to inflammation and fibrosis. KEGG pathway analysis of DEGs revealed that the major enriched pathways were related to protein digestion and absorption, diabetic cardiomyopathy, AGE-RAGE signaling in diabetes, proteoglycans in cancer, and ECM-receptor interaction.

The GO biological pathways enrichment analysis also revealed that the major enriched biological pathways in WT mice after TAC were related to collagen fibril organization, extracellular matrix organization, cell adhesion, ossification, angiogenesis, and TGF-β signaling. The findings that IL-12β KO significantly suppressed the above LV gene profiles, particularly the gene profiles associated with extracellular collagen formation and matrix remodeling and inflammation, demonstrate that IL-12β exerts important and broad roles in promoting systolic overload-induced LV inflammation, fibrosis, and dysfunction. The drastic reductions of multiple LV signaling pathways of extracellular matrix remodeling and immune related pathways in IL-12β KO mice after TAC not only indicate a critical role of IL-12β in promoting LV fibrosis and inflammation but also support the notion that there is significant crosstalk/interaction between extracellular matrix remodeling and immune cells during TAC-induced LV inflammation and dysfunction.

Several findings from this study also suggest that T cells, particular CD8^+^ T cells, might play an important role in the reduced LV inflammation and dysfunction in IL-12β KO mice after TAC: (i) Both LV CD8^+^ cell activation and CD4^+^ cell activation were reduced in IL-12β KO mice after TAC; (ii) LV IFNγ^+^CD8^+^ cells in cardiac drainage lymph nodes were significantly reduced in IL-12β KO mice after TAC; (iii) IL-12β KO significantly reduced TAC-induced expression of multiple adhesion molecules (such as Cxadr, CD44, Icam1, and CD29) for recruiting immune cells and beyond. Cxadr is a protein that modulates both MHCI molecule processing and immune cell infiltration; (iv) Expressions of CXCR3 and CD44 in CD8^+^ T cells and CD4^+^ T cells in drainage lymph nodes were significantly reduced in IL-12β KO as compared with WT mice after TAC. (v) IL-12β KO significantly decreased expression of LV genes that could promote CD8 T cells’ effective function (such as B2m, MHCI molecules). Thus, these findings support that IL-12β KO protects hearts against TAC-induced inflammation at least partially by restraining the overactive immune cells, particularly CD8^+^ T cells. While the reduced cardiac inflammation in IL-12β KO is likely a collective effect of multiple immune cells and nonimmune cells, the reduced IL-12 and IL-23 by IL-12β KO likely play a central role in protecting hearts against TAC-induced LV inflammation and HF.

This study has several limitations. First, we found that IL-12β KO attenuated TAC-induced cardiac inflammation, cardiomyocyte hypertrophy, and fibrosis. As each of these above factors could independently contribute to HF development, our study is unable to fully determine the relative role of IL-12β in modulating LV inflammation, cardiomyocyte hypertrophy, and fibrosis. Second, HF could progress to lung inflammation and remodeling, as well as RV hypertrophy, while lung inflammation can also directly promote lung remodeling and RV hypertrophy. Thus, we could not identify whether the reduced lung inflammation and remodeling is partially due to the direct impact on lung inflammation. Nevertheless, this study underscores an important role for IL-12β inhibition to attenuate pressure overload-induced LV inflammation and HF development.

In summary, our data clearly demonstrate that IL-12β plays an important role in promoting pressure overload-induced cardiac dysfunction, hypertrophy, inflammation, fibrosis, and the consequent pulmonary inflammation and remodeling, and RV hypertrophy. These findings suggest that inhibition of IL-12β signaling may be a viable approach to treat systolic overload-induced cardiac inflammation and dysfunction.

## Acknowledgment

The authors would like to acknowledge the Histology and Flow Cytometry Core Facility of the Department of Physiology and Biophysics at the University of Mississippi Medical Center.

