## Supplementary for "Genetic inhibition of IL-12β suppresses systolic overload-induced cardiac inflammation and heart failure development"

**Supplementary Materials**

**Histological Staining:** Cardiac and pulmonary samples were fixed in 4% formaldehyde, embedded in paraffin blocks, and sections of 5 µm were sliced. The tissue sections were deparaffinized and rehydrated. Sirius Red/Fast Green Staining Kit (Chondrex Inc., 90461) was used for LV and lung fibrosis staining. Alexa Flour-488 conjugated wheat germ agglutinin (WGA) (Invitrogen, W11261, 5 µg/mL) staining was used to measure LV cardiomyocyte cross-sectional area. The cross-sectional area of 100 LV cardiomyocytes was measured and averaged to get the mean cardiomyocyte cross-sectional area. Fibrosis and LV cardiomyocyte cross-sectional area were quantified using ImageJ software from the National Institutes of Health. The infiltrated CD45^+^ leukocytes in the LV and lung tissues were stained with goat anti-CD45 antibody (R&D Systems, AF114, 1:100 dilution). The infiltrated CD45^+^ leukocytes were visualized using Alexa Flour-555 conjugated donkey anti-goat secondary antibody (Invitrogen, A21432, 1:1000 dilution). Muscularization of pulmonary arterioles was determined by using mouse anti-α smooth muscle actin (αSMA) (Invitrogen, 14-9760-82, 1:200 dilution) and rabbit anti-CD31 antibody (Cell Signaling Technologies Inc., 77699, 1:200 dilution). The αSMA and CD31 were then visualized by using Alexa Fluor-555 conjugated goat anti-mouse (Invitrogen, A21424, 1:1000 dilution) and Alexa Fluor-594 conjugated goat anti-rabbit (Invitrogen, A11012, 1:1000 dilution) secondary antibodies, respectively. The tissue sections were then mounted with 4',6-diamidino-2-phenylindole (DAPI) containing mounting media (enQuire BioReagents, QS4-20ML) to visualize the nucleus. All the histological images were captured using Mantra Quantitative Pathology Imaging System (Perkin Elmer), and infiltrated CD45^+^ leukocytes were quantified using inForm software version 2.2.1 (Perkin Elmer).

**Western Blot Analyses:** LV tissues were homogenized in RIPA lysis buffer supplied with 1X protease and phosphatase inhibitor cocktail. The protein content of LV tissue lysate was determined by using the BCA protein assay kit. Then, 50µg LV tissue lysate was loaded into each well, separated by SDS-PAGE, and transferred into a methanol-activated PVDF membrane at 100 mV for 100 minutes. The membranes were then blocked with 5% non-fat milk in TBST for 1 hour at room temperature in a rocker and incubated with primary antibodies for IL-12β (BioXCell, Catalog # BE0051, 1:1000 dilution) and β-actin (Cell Signaling Technologies Inc., Catalog # 4967, 1:1000 dilution) overnight at 4°C in a rocker. The membranes were then washed with TBST 3 times, 10 minutes each, and were incubated with HRP-conjugated secondary antibodies (Abcam, 1:4000 dilution in 5% non-fat dry milk in TBST) for 1 hour at room temperature in a rocker. Then, the protein bands were detected using the iBright FL1500 instrument (ThermoFisher Scientific). The relative protein expression was quantified using ImageJ software from NIH.

**Supplementary Table 1:** Antibodies used for flow cytometry analyses

| **Antibody** | **Conjugate** | **Clone** | **Vendor** | **Catalog #** |
| --- | --- | --- | --- | --- |
| **CD3e** | BUV737 | 145-2C11 | BD Biosciences | 612771 |
| **CD4** | BUV496 | GK1.5 | BD Biosciences | 612952 |
| **CD8α** | BB790 | 53-6.7 | BD Biosciences | 624296 |
| **CD11b** | BV650 | M1/70 | Biolegend | 101259 |
| **CD11c** | BV711 | N418 | Biolegend | 117349 |
| **CD16/32** | - | 93 | Biolegend | 101302 |
| **CD19** | BUV395 | 1D3 | BD Biosciences | 563557 |
| **CD44** | FITC | IM-7 | BD Biosciences | 553133 |
| **CD45** | BUV805 | 30-F11 | BD Biosciences | 568336 |
| **CD62L** | AF700 | MEL-14 | BD Biosciences | 104418 |
| **F4/80** | BUV563 | T45-2342 | BD Biosciences | 749284 |
| **I-A/I-E (MHC-II)** | APC-Cy7 | M5/114.15.2 | Biolegend | 107628 |
| **Ly6C** | BV605 | AL-21 | BD Biosciences | 563011 |
| **Ly6G** | AF700 | 1A8 | Biolegend | 127622 |
| **IFNγ** | FITC | XMG1.2 | BioLegend | 505806 |
| **PD1** | BV480 | 29F.1A12 | BD Biosciences | 568612 |
| **CXCR6** | BV711 | SA051D1 | BioLegend | 151111 |
| **IL17A** | APC-Cy7 | TC11-18H10.1 | BioLegend | 506940 |
| **Pro-IL1β** | PE-Cy7 | NJTEN3 | Invitrogen | 25-7114-82 |
| **IL10** | PE | JES5-16E3 | Invitrogen | 12-7101-82 |

**Supplementary Table 2.** Anatomic data of male wild-type (WT) and IL-12β knockout (IL-12β KO) mice under sham and TAC conditions

| **Parameters** | **WT Sham**  **(n=9)** | **IL-12β KO Sham**  **(n=18)** | **WT TAC**  **(n=10)** | **IL-12β KO TAC (n=14)** |
| --- | --- | --- | --- | --- |
| Body Weight (g) | 30.5±0.41 | 29.34±0.53 | 31.61±0.65 | 28±0.53**^†^** |
| Tibial Length (mm) | 18.17±0.11 | 18.05±0.056 | 18.42±0.06 | 17.9±0.09 |
| LV Weight (mg) | 106.3±2.16 | 98.11±1.87 | 197.96±9.62* | 133.6±4.71^#^**^†^** |
| LA Weight (mg) | 4.14±0.35 | 3.59±0.14 | 12.43±1.88* | 5.0±0.43**^†^** |
| Lung Weight (mg) | 147.8±5.1 | 146.48±2.4 | 290.8±34.78* | 143.6±4.03**^†^** |
| RV Weight (mg) | 21.41±0.41 | 21.22±0.79 | 30.17±2.57* | 19.3±0.7**^†^** |
| RA Weight (mg) | 3.67±0.34 | 3.61±0.18 | 4.64±0.44 | 3.6±0.24 |
| Total Heart Weight (mg) | 135.53±2.94 | 126.53±2.51 | 245.2±13.73* | 161.5±5.44^#^ |
| LV weight/BW (mg/g) | 3.49±0.06 | 3.35±0.06 | 6.29±0.35* | 4.8±0.19**^†^** |
| LA weight/BW (mg/g) | 0.14±0.01 | 0.12±0.005 | 0.4±0.07* | 0.2±0.02**^†^** |
| Lung weight/BW (mg/g) | 4.84±0.14 | 5±0.09 | 9.32±1.26* | 5.1±0.13**^†^** |
| RV weight/BW (mg/g) | 0.7±0.01 | 0.72±0.02 | 0.97±0.1* | 0.7±0.02**^†^** |
| RA weight/BW (mg/g) | 0.12±0.01 | 0.12±0.006 | 0.15±0.02 | 0.1±0.007 |

Data are mean ± SEM. *p<0.05 compared with WT Sham, #p<0.05 compared with IL-12β KO Sham, **^†^**p<0.05 compared with WT TAC; BW, body weight.

**Supplementary Table 3.** Anatomic data of female wild-type (WT) and IL-12β knockout (IL-12β KO) mice under sham and TAC conditions

| **Parameters** | **WT Sham**  **(n=7)** | **IL-12β KO Sham**  **(n=14)** | **WT TAC**  **(n=16)** | **IL-12β KO TAC (n=22)** |
| --- | --- | --- | --- | --- |
| Body Weight (g) | 25.3±0.61 | 24.66±0.71 | 21.11±0.78* | 24.1±0.5**^†^** |
| Tibial Length (mm) | 17.53±0.08 | 17.62±0.11 | 17.35±0.11 | 17.5±0.08 |
| LV Weight (mg) | 79.74±2.65 | 80.69±1.96 | 149.57±4.55* | 128.2±4.04^#^**^†^** |
| LA Weight (mg) | 2.61±0.14 | 2.84±0.23 | 13.6±2.65* | 6.5±0.67**^†^** |
| Lung Weight (mg) | 144.54±4.67 | 144.54±4.38 | 268.08±33.17* | 163.4±8.15**^†^** |
| RV Weight (mg) | 15.69±0.62 | 16.41±0.61 | 24.69±1.35* | 19±0.57^#^**^†^** |
| RA Weight (mg) | 2.87±0.08 | 3.14±0.24 | 4.83±0.39* | 3.4±0.13**^†^** |
| Total Heart Weight (mg) | 100.91±2.86 | 103.09±2.4 | 192.69±8.1* | 157.1±5.03^#^**^†^** |
| LV weight/BW (mg/g) | 3.15±0.05 | 3.29±0.08 | 7.31±0.45* | 5.4±0.16^#^**^†^** |
| LA weight/BW (mg/g) | 0.1±0.005 | 0.12±0.007 | 0.71±0.16* | 0.3±0.03**^†^** |
| Lung weight/BW (mg/g) | 5.71±0.06 | 5.89±0.16 | 13.62±2.07* | 6.9±0.37**^†^** |
| RV weight/BW (mg/g) | 0.62±0.03 | 0.67±0.02 | 1.23±0.11* | 0.8±0.03**^†^** |
| RA weight/BW (mg/g) | 0.11±0.004 | 0.13±0.009 | 0.24±0.03* | 0.1±0.006**^†^** |

Data are mean ± SEM. *p<0.05 compared with WT Sham, #p<0.05 compared with IL-12β KO Sham, **^†^**p<0.05 compared with WT TAC; BW, body weight.

**Supplementary Table 4.** Anatomic data of male wild-type (WT) and IL-12β knockout (IL-12β KO) mice one week after TAC

| **Parameters** | **WT TAC**  **(n=8)** | **IL-12β KO TAC**  **(n=10)** |
| --- | --- | --- |
| Body Weight (g) | 28.35±0.83 | 25.6±1.08 |
| Tibial Length (mm) | 17.38±0.04 | 17±0.17 |
| LV Weight (mg) | 144.21±7.93 | 116.9±4.52* |
| LA Weight (mg) | 6.88±0.71 | 5.1±0.5* |
| Lung Weight (mg) | 232.26±16.05 | 188±13.35* |
| RV Weight (mg) | 20.83±0.58 | 19.6±0.32 |
| RA Weight (mg) | 3.84±0.51 | 3.4±0.16 |
| Total Heart Weight (mg) | 175.75±8.96 | 144.9±5.19* |
| LV weight/BW (mg/g) | 5.09±0.25 | 4.6±0.2 |
| LA weight/BW (mg/g) | 0.24±0.02 | 0.2±0.02 |
| Lung weight/BW (mg/g) | 8.27±0.67 | 7.5±0.64 |
| RV weight/BW (mg/g) | 0.74±0.02 | 0.8±0.02 |
| RA weight/BW (mg/g) | 0.13±0.009 | 0.1±0.008 |

Data are mean ± SEM. *p<0.05 compared to WT TAC; BW, body weight.

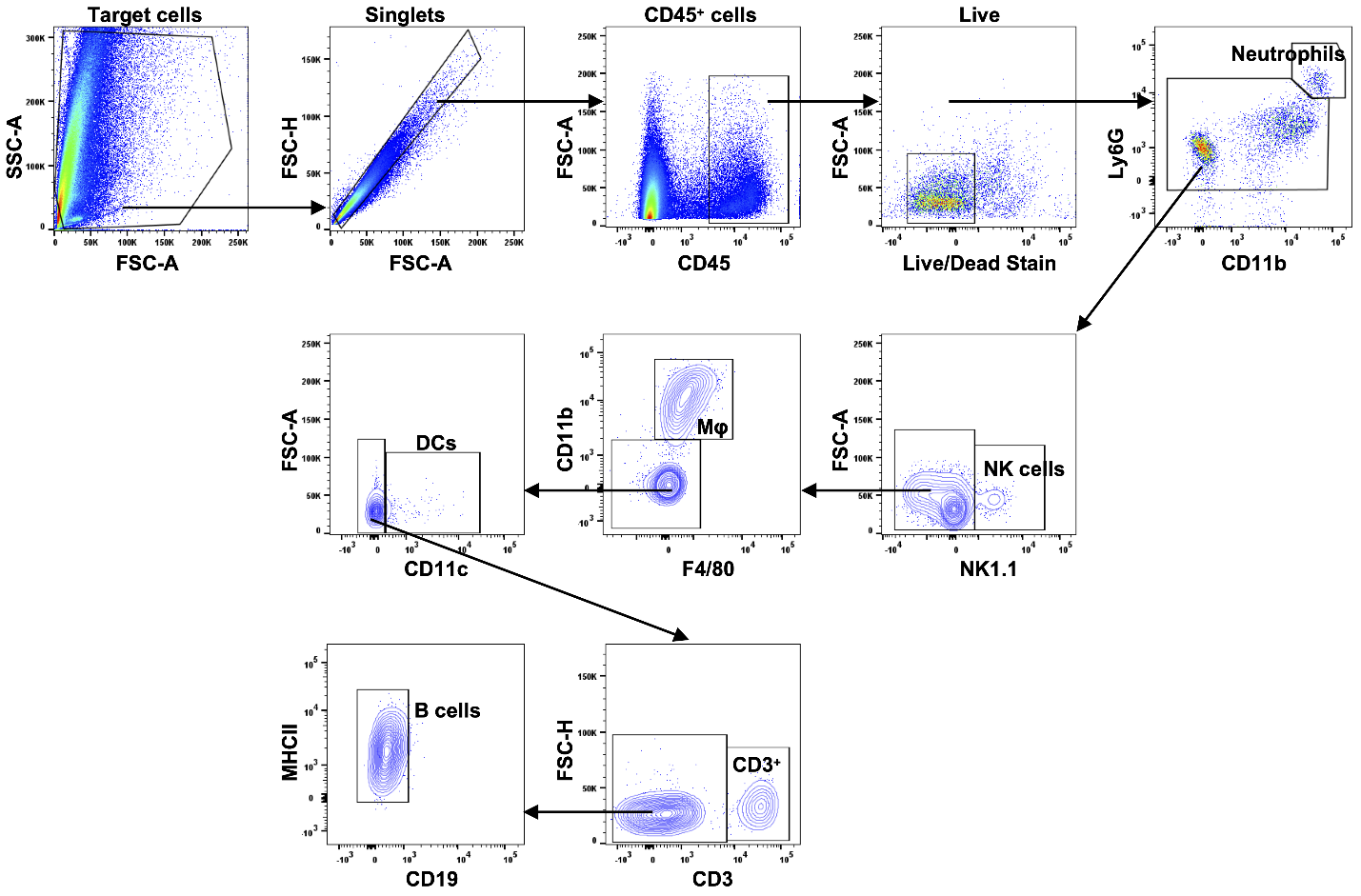

**Supplementary Figure 1:** Flow cytometry gating strategy used for identification of Neutrophils, NK cells, Macrophages, Dendritic cells, T cells, and B cells in LV tissue; Mφ, Macrophages; DCs, Dendritic cells.

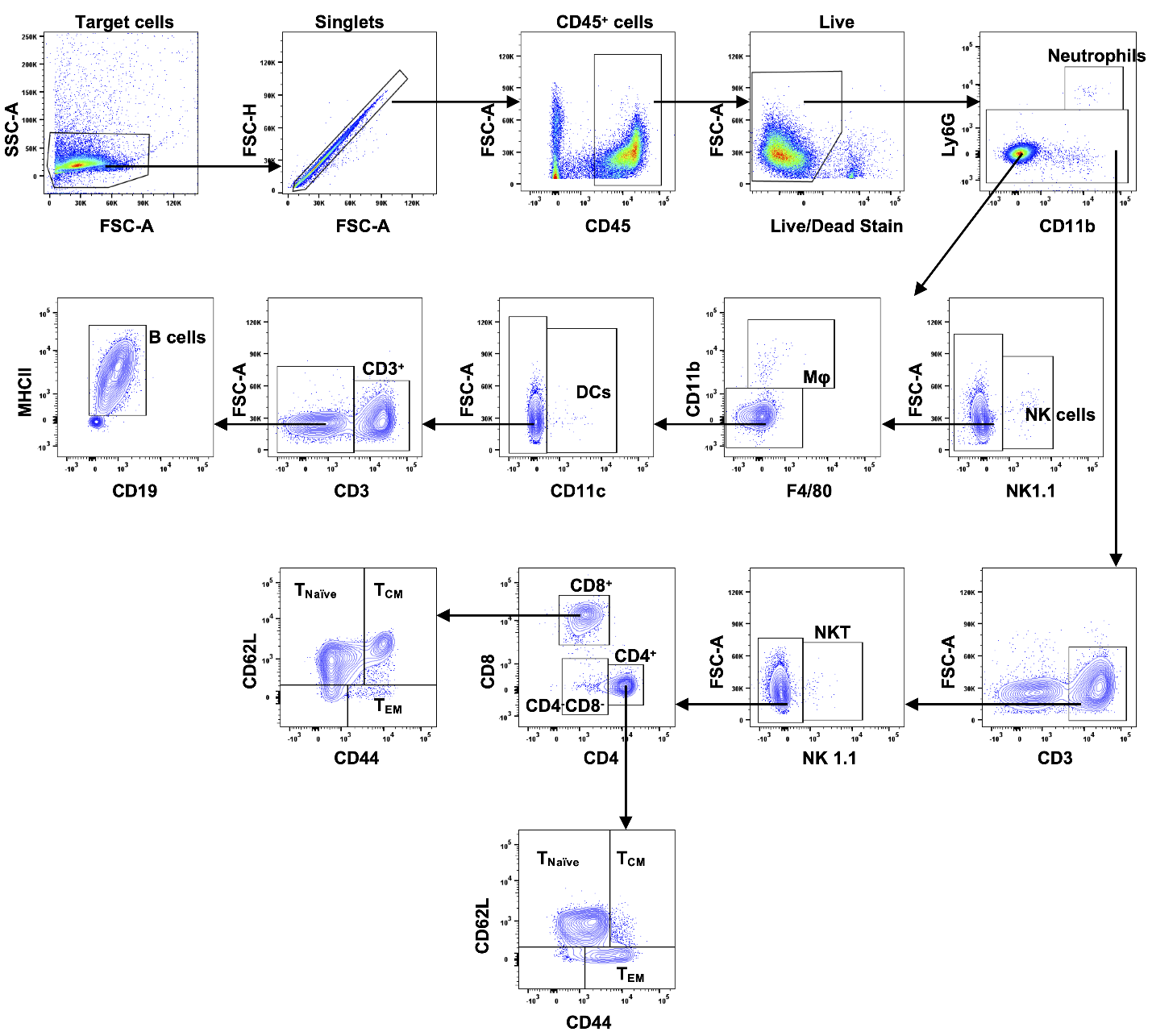

**Supplementary Figure 2:** Flow cytometry gating strategy used for identification of Neutrophils, NK cells, Macrophages, Dendritic cells, T cells, and B cells in cardiac drainage lymph node; Mφ, Macrophages; DCs, Dendritic cells; T_EM_, Effective Memory T Cells; T_naïve_, Naïve T Cells; T_CM_, Central Memory T Cells.

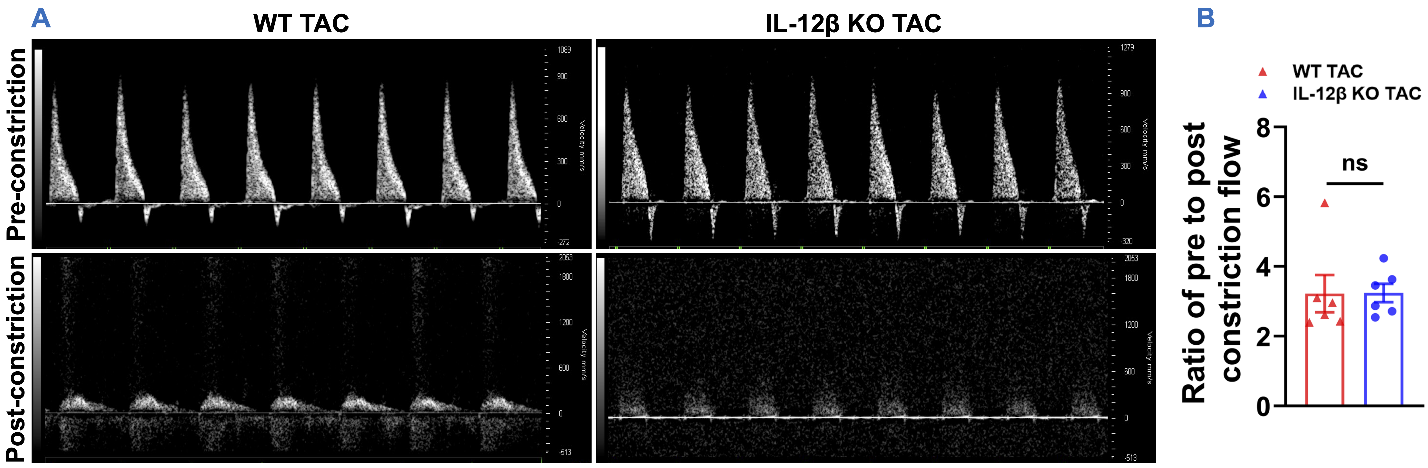

**Supplementary Figure 3:** (A) Representative pulsed wave Doppler images of pre- and post-constriction sites of WT and IL-12β KO mice after TAC. (B) Quantified data of the flow difference across the constriction site.

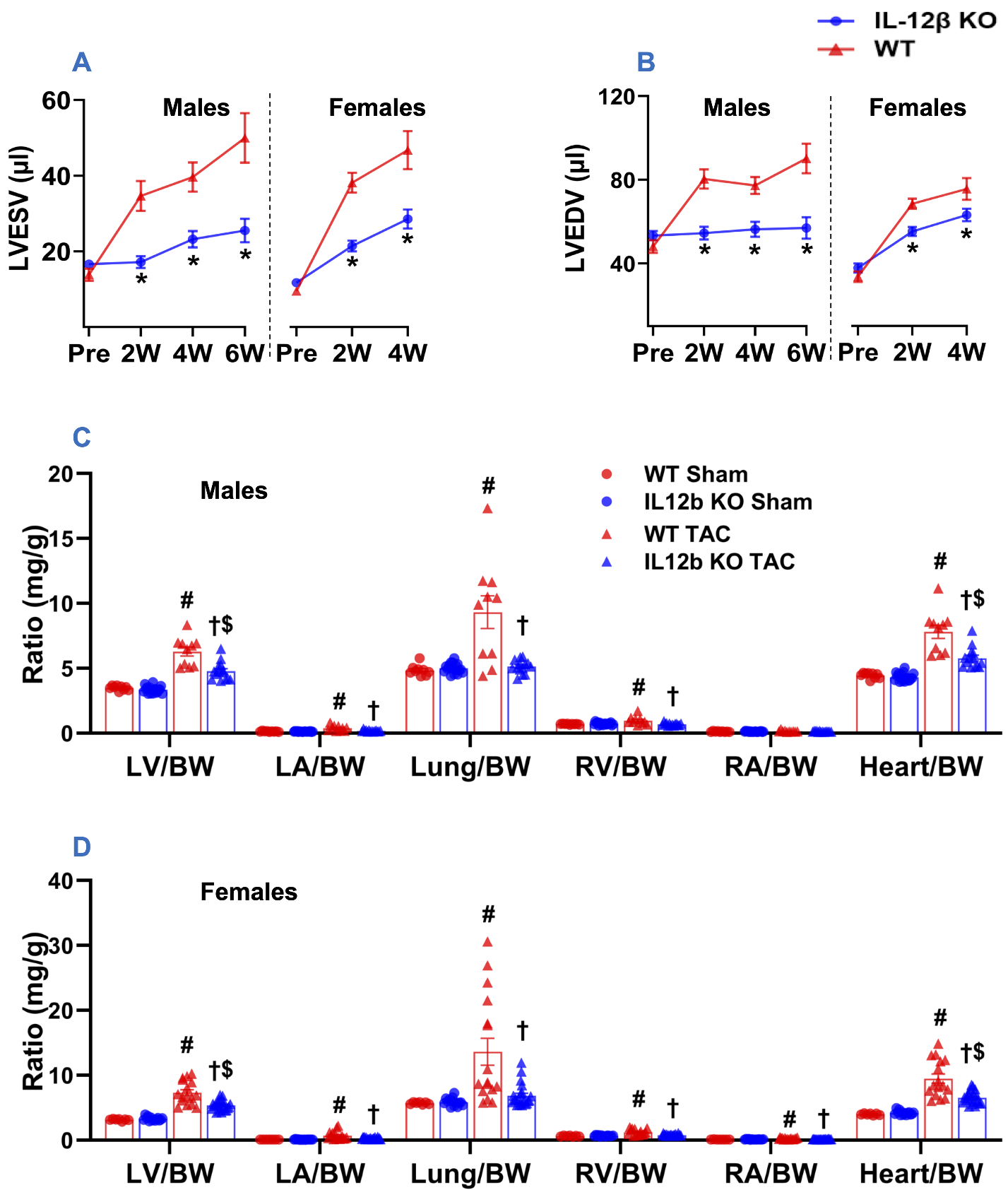

**Supplementary Figure 4:** (A, B) Quantified data of echocardiographic measurements of LV end-systolic and end-diastolic volume of both sexes. (C, D) The ratio of LV weight, left atrial (LA) weight, lung weight, RV weight, right atrial (RA) weight, and total heart weight to body weight (BW) of the indicated groups. *p<0.05; ^#^p<0.05 compared to WT sham; ^†^p<0.05 compared to WT TAC; ^$^p<0.05 compared to IL-12β KO sham; n = 7 to 22 per group; LVESV, LV end-systolic volume; LVEDV, LV end-diastolic volume.

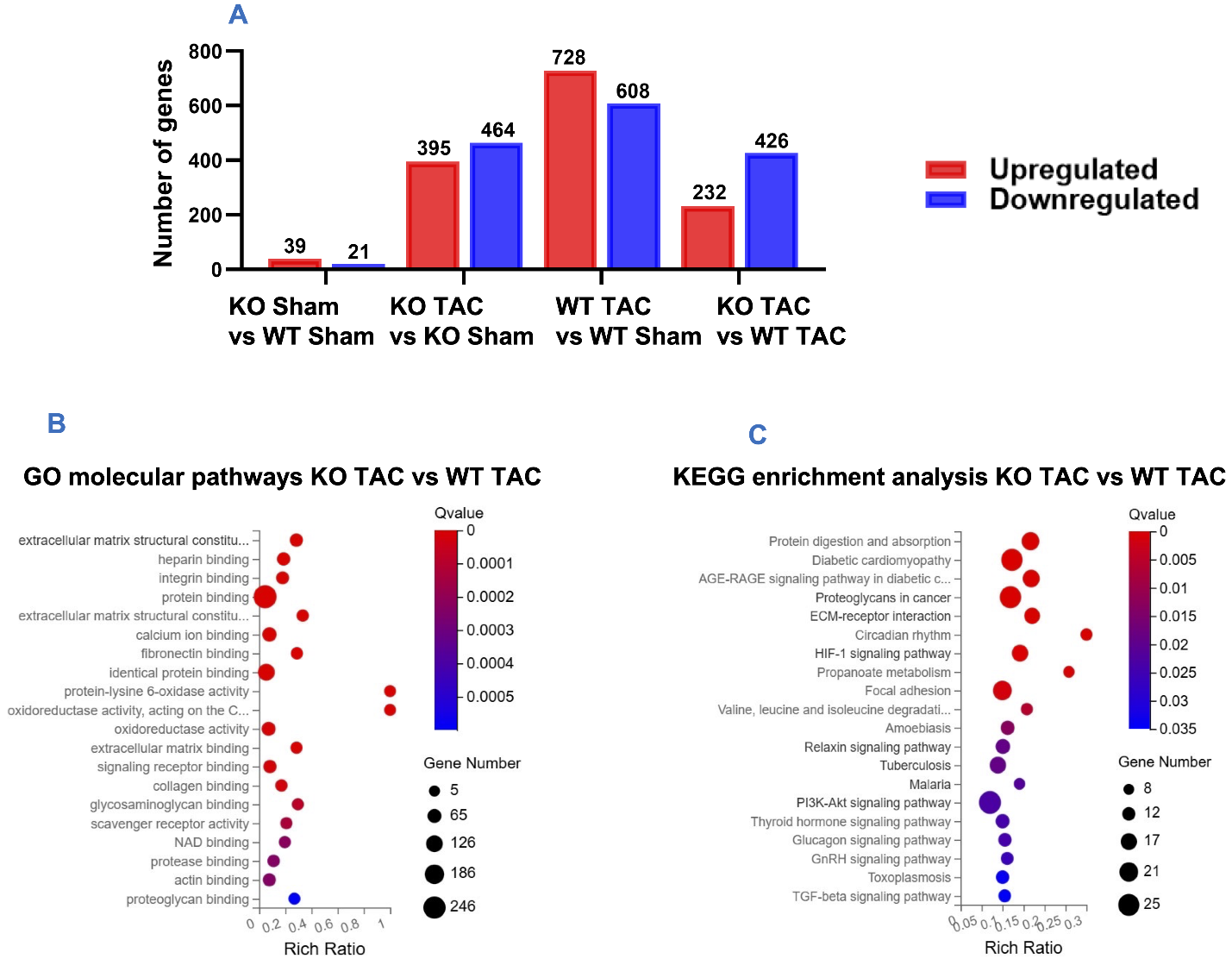

**Supplementary Figure 5:** (A) Statistics of differentially expressed genes (DEGs) of the indicated groups. (B) Top enriched GO molecular pathways of KO TAC vs WT TAC. (C) Top enriched KEGG pathways of KO TAC vs WT TAC. n=2-3 per group.

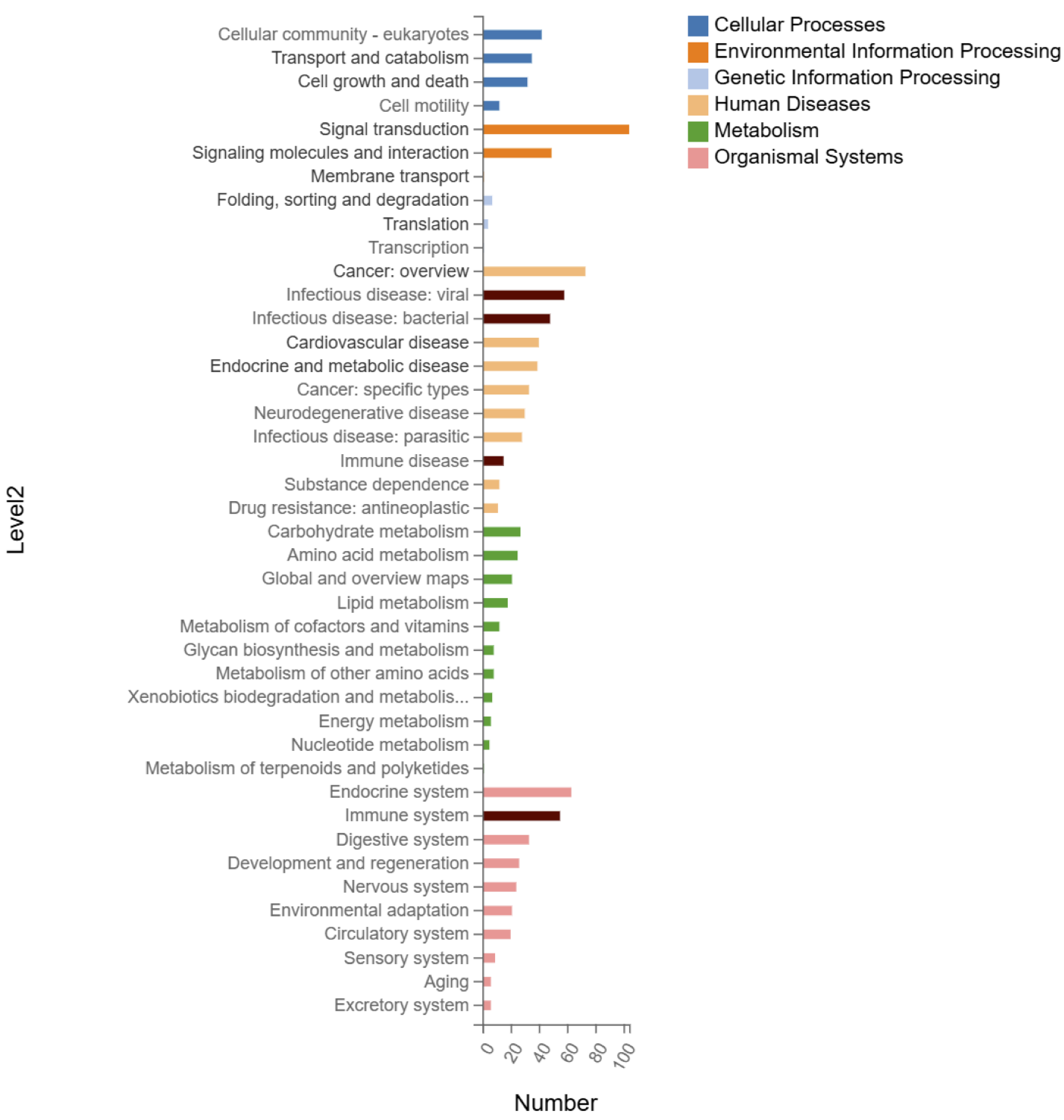
**Supplementary Figure 6:** KEGG pathway classification of WT TAC vs KO TAC mice.

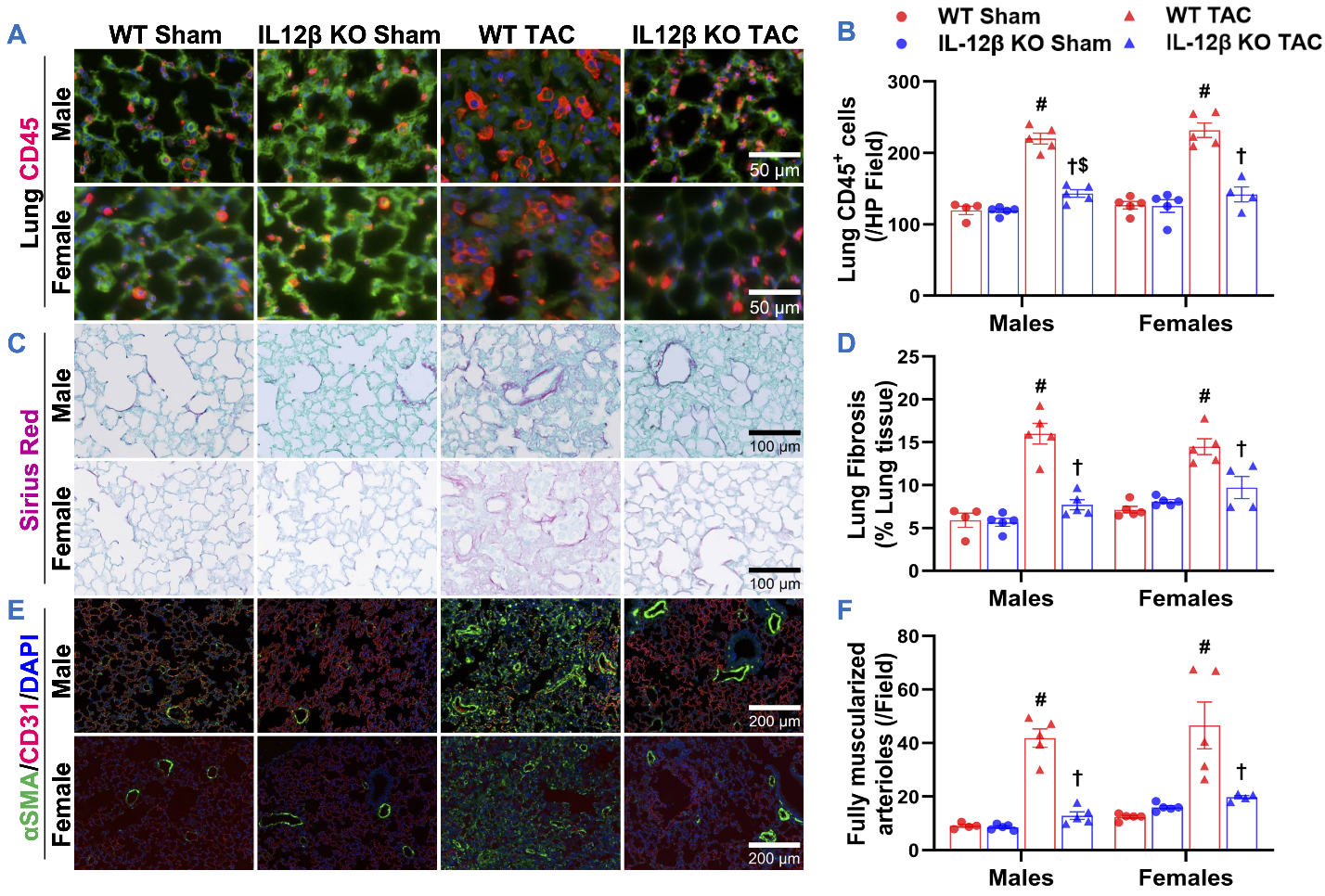

**Supplementary Figure 7: IL-12β KO attenuated TAC-induced pulmonary inflammation, fibrosis, and vessel remodeling in male and female mice.** (A, B) Representative images and quantified data of infiltrated CD45^+^ leukocytes in the lung performed by immuno-histological staining. (C, D) Representative images and quantified data of lung fibrosis performed by Sirius Red/Fast Green staining. (E, F) Representative images and quantified data of lung vessel remodeling performed by immuno-histological staining. ^#^p<0.05 compared to WT sham; ^†^p<0.05 compared to WT TAC; ^$^p<0.05 compared to IL-12β KO sham; n = 4-5 per group.

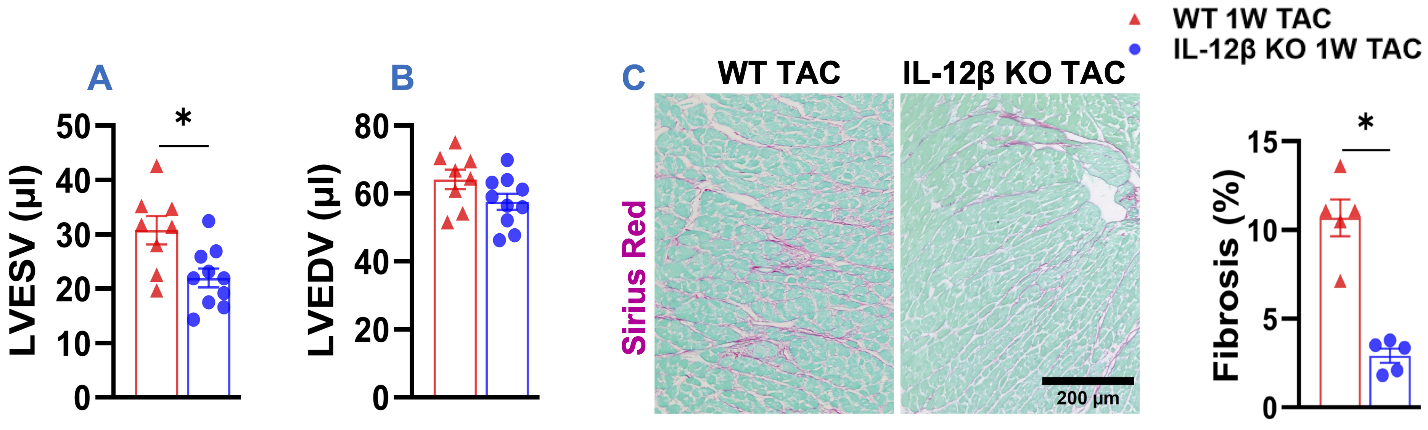

**Supplementary Figure 8: IL-12β KO significantly attenuated TAC-induced early-phase LV dysfunction and fibrosis.** (A, B) Quantified data of echocardiographic measurements of LV end-systolic volume (LVESV) and LV end-diastolic volume (LVEDV) of the indicated groups, respectively. (C) Representative images and quantified data of LV fibrosis of the indicated groups. *p<0.05; n = 5-10 per group.

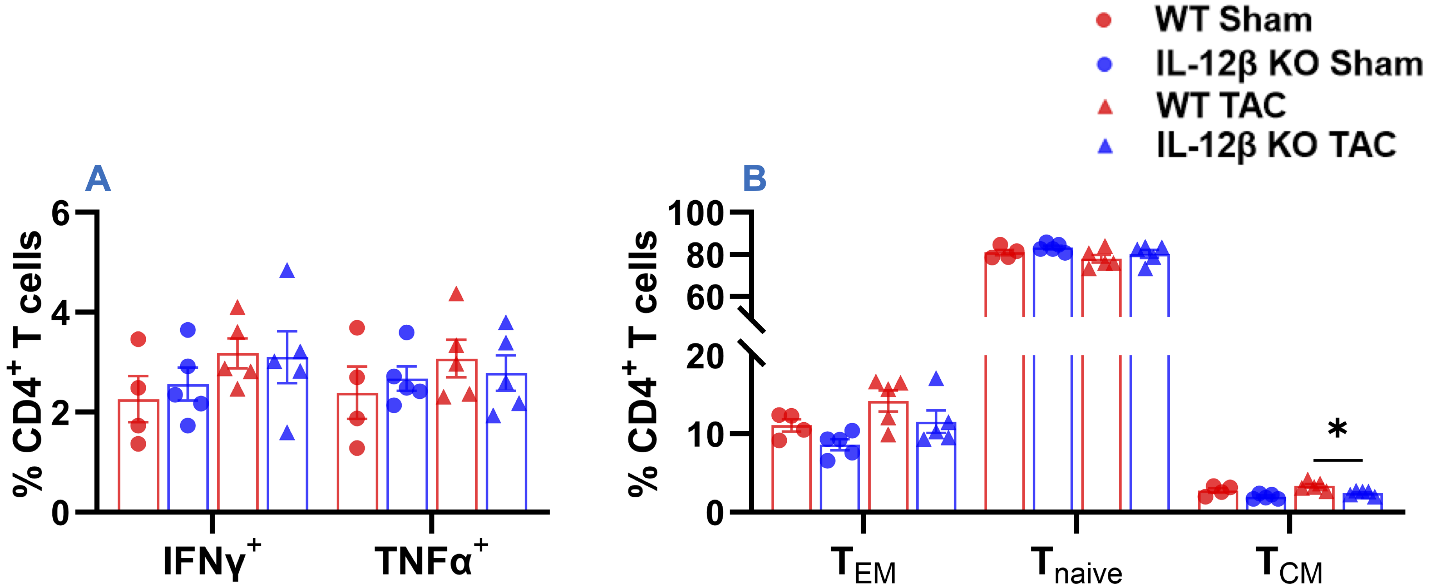

**Supplementary Figure 9:** (A) Percentages of IFNγ^+^CD4^+^ and TNFα^+^CD4^+^ within CD4^+^ T cells. (B) Percentage of effective memory (CD44^+^CD62L^-^), naïve (CD44^-^CD62L^+^), and central memory (CD44^+^CD62L^+^) T cells of CD4^+^ T cells. *p<0.05; T_EM_, Effective Memory T Cells; T_naïve_, Naïve T Cells; T_CM_, Central Memory T Cells; n = 5 per group.
